# Pharmaceutical-grade Rigosertib is a Microtubule-destabilizing Agent

**DOI:** 10.1101/2020.01.28.923235

**Authors:** Marco Jost, Yuwen Chen, Luke A. Gilbert, Max A. Horlbeck, Lenno Krenning, Grégory Menchon, Ankit Rai, Min Y. Cho, Jacob J. Stern, Andrea E. Prota, Martin Kampmann, Anna Akhmanova, Michel O. Steinmetz, Marvin E. Tanenbaum, Jonathan S. Weissman

**Author notes:** Correspondence (M.E.T.), (J.S.W.).

## Abstract

We recently used CRISPRi/a-based chemical-genetic screens and targeted cell biological, biochemical, and structural assays to determine that rigosertib, an anti-cancer agent in phase III clinical trials, kills cancer cells by destabilizing microtubules. In a recent manuscript, Reddy and co-workers suggest that this microtubule-destabilizing activity of rigosertib is mediated not by rigosertib itself but by a contaminating degradation product of rigosertib, ON01500, present in formulations obtained from commercial vendors (Baker et al., 2019). Here, we demonstrate that treatment of cells with pharmaceutical-grade rigosertib (>99.9% purity) results in qualitatively indistinguishable phenotypes as treatment with commercially obtained rigosertib across multiple assays. The two compounds have indistinguishable chemical-genetic interactions with genes involved in modulating the microtubule network (*KIF2C* and *TACC3*), both destabilize microtubules in cells and *in vitro*, and both show substantially reduced toxicity in cell lines expressing a rationally-designed mutant of tubulin (L240F *TUBB* mutant), in which the rigosertib binding site in tubulin is mutated. Importantly, the specificity of the L240F *TUBB* mutant for microtubule-destabilizing agents, which is disputed by Reddy and co-workers, was recently confirmed by an independent research group (Patterson et al., 2019). We conclude that rigosertib kills cancer cells by destabilizing microtubules, in agreement with our original findings.

## Introduction

The case of rigosertib is a classic example of pleiotropic effects confounding targeted assays; depending on the type of assay, supposed evidence has emerged for multiple conflicting molecular targets. It is worth outlining the history of rigosertib’s development here to illustrate this issue. Rigosertib was first described by Reddy and co-workers in 2005 as an in vitro inhibitor of polo-like kinase 1 (PLK1) and proposed to kill cancer cells through this activity, based on measurements of cell cycle progression and cellular PLK1 activity (Gumireddy et al., 2005). This claim was disputed by Steegmaier et al. in 2007, who found that the cellular phenotypes induced by rigosertib did not match those induced by the *bona fide* PLK1 inhibitor BI2536, with rigosertib’s suppression of cellular PLK1 activity likely being an indirect effect (Steegmaier et al., 2007). A later study using a FRET sensor for PLK1 activity in cells similarly found no evidence for PLK1 inhibition by the compound (Mäki-Jouppila et al., 2014). Rigosertib was then proposed to target PI3 kinase by Reddy and co-workers as well as others, based on inhibition of PI3 kinase signaling in rigosertib-treated cells (Prasad et al., 2009; Chapman et al., 2012; Hyoda et al., 2015), but subsequent work by others could not confirm the direct inhibition of PI3 kinase (Mäki-Jouppila et al., 2014). Presenting data from in vitro binding assays and measurements of phosphorylation state of proteins in the RAS signaling cascade, Reddy and co-workers then proposed in 2016 that rigosertib directly inhibits RAS signaling by engaging RAS-binding domains of effector proteins and preventing interaction of these effectors with RAS (Athuluri-Divakar et al., 2016). This proposal, however, was refuted by Ritt et al. later in 2016 who found that rigosertib did not appreciably block interaction of RAS with the RAS-binding domain of RAF but that rigosertib instead, either directly or indirectly, activates JNK signaling, leading to hyperphosphorylation of several RAS effectors including RAFs and SOS1 and thereby inhibiting RAS signaling (Ritt et al., 2016). Thus, Ritt et al. concluded that the effect of rigosertib on RAS signaling was indirect, with the actual molecular target still left open. Intriguingly, a large-scale microscopy-based screen revealed a striking phenotypic similarity between rigosertib and microtubule-targeting agents, pointing towards microtubules as a possible target for rigosertib (Twarog et al., 2016). Despite the uncertainty over its mechanism, rigosertib progressed through clinical trials and at the time of our initial study was in phase III clinical trials for myelodysplastic syndrome and earlier stage trials for several other cancers. Thus far, however, further progression is hampered by a lack of efficacy in the general patient population (Garcia-Manero et al., 2016; O’Neil et al., 2015).

In light of this ambiguity, we considered rigosertib to be an excellent test case for unbiased genetic approaches that explore the full spectrum of all possible mechanisms simultaneously. We therefore developed a strategy based on combined genome-wide CRISPR-based knockdown and overexpression screens to probe rigosertib’s genetic dependencies systematically (Jost et al., 2017). These screens revealed that destabilization of microtubules, for example by overexpression of the microtubule depolymerase MCAK (encoded by *KIF2C*) or knockdown of the microtubule-stabilizing factor *TACC3*, sensitized cells to rigosertib, whereas stabilization of microtubules protected cells against rigosertib, suggesting that rigosertib might be a microtubule-destabilizing agent. Indeed, subsequent targeted assays confirmed that rigosertib directly inhibits microtubule polymerization in cells and in vitro, and a co-crystal structure of rigosertib bound to tubulin revealed that rigosertib binds in the colchicine site of β-tubulin. Guided by the structure, we designed a point mutation in β-tubulin to abrogate rigosertib binding (L240F *TUBB*). Expression of this mutant conferred resistance to rigosertib in three different cell lines. Critically, the resistance was specific to agents with the same binding mode as rigosertib but not vinblastine, a microtubule-destabilizing agent that binds to a different site on tubulin. In a recent manuscript, Patterson et al. report that the MTH1 inhibitor TH588 also destabilizes microtubules by binding to the same site as rigosertib, as evidenced by a co-crystal structure (Patterson et al., 2019). Patterson et al. found that our L240F *TUBB* mutant provided resistance against TH588 but not against the PLK1 inhibitor BI2536, further confirming the specificity of the resistance conferred by the L240F mutant (Patterson et al., 2019). Together, our results strongly suggested that rigosertib kills cancer cells by directly destabilizing microtubules.

In a recent manuscript, Reddy and co-workers argue that rigosertib does not have microtubule-destabilizing activity (Baker et al., 2019). They instead suggest that the microtubule-destabilizing activity is mediated by ON01500, a product of photodecarboxylative degradation of rigosertib that is present in commercially available rigosertib, but not in pharmaceutical-grade rigosertib. We have now obtained pharmaceutical-grade rigosertib from Onconova (the company that supplies rigosertib for clinical trials) and demonstrate using multiple assays that pharmaceutical-grade rigosertib also directly destabilizes microtubules and kills cells through this microtubule-destabilizing activity, fully consistent with our original findings.

## Results

We conducted a series of assays with pharmaceutical-grade rigosertib (rigosertib_pharm_) obtained from Onconova, mirroring the assays in our original study. Where indicated, control experiments were also performed with pure ON01500 (Onconova) and commercially obtained rigosertib (rigosertib_comm_). Throughout our experiments, we took precautions to prevent pH- or light-induced degradation of rigosertib. We minimized light exposure by keeping the lights in our tissue culture hoods off during work with rigosertib and by minimizing the duration any rigosertib-treated cultures spent outside of the incubators. Furthermore, we prepared our rigosertib stocks as instructed by scientists at Onconova and by Dr. Reddy. Specifically, for all experiments shown in this manuscript we prepared rigosertib stocks freshly by dissolving solid directly in DMSO and made dilutions in PBS to prevent pH drops. For imaging experiments, we used the lowest possible laser power and exposure times (100 ms or less) to minimize light exposure.

### Pharmaceutical-grade and commercial rigosertib have identical chemical-genetic interactions

We first assessed how genetic destabilization of microtubules affects sensitivity to rigosertib_pharm_, rigosertib_comm_, and pure ON01500. Specifically, we used internally-controlled drug sensitivity assays to measure how drug sensitivity is affected by knockdown or overexpression of *KIF2C* and *TACC3*, both of which modulate microtubule stability (see above). We transduced K562 CRISPRi or CRISPRa cells with BFP-marked sgRNA expression vectors at MOI < 1 (15-40% of the population expressed sgRNAs) and then tracked the fraction of sgRNA-expressing (BFP-positive) cells in the population after treatment with the drugs or with DMSO (control) to determine how expression of each sgRNA affects growth in the presence of the different drugs. In agreement with our original findings, cells were sensitized to rigosertib_pharm_ by either knockdown of *TACC3* or overexpression of *KIF2C* (Fig. 1A), and vice versa overexpression of *TACC3* or knockdown of *KIF2C* protected against rigosertib_pharm_. We observed the same drug sensitivity phenotypes for both ON01500 and rigosertib_comm_, although we note that ON01500 was substantially more toxic and rigosertib_comm_ was slightly more toxic than rigosertib_pharm_, giving rise to variable selective pressures. These results establish that genetic destabilization of microtubules also sensitizes to rigosertib_pharm_.

**Figure 1.**
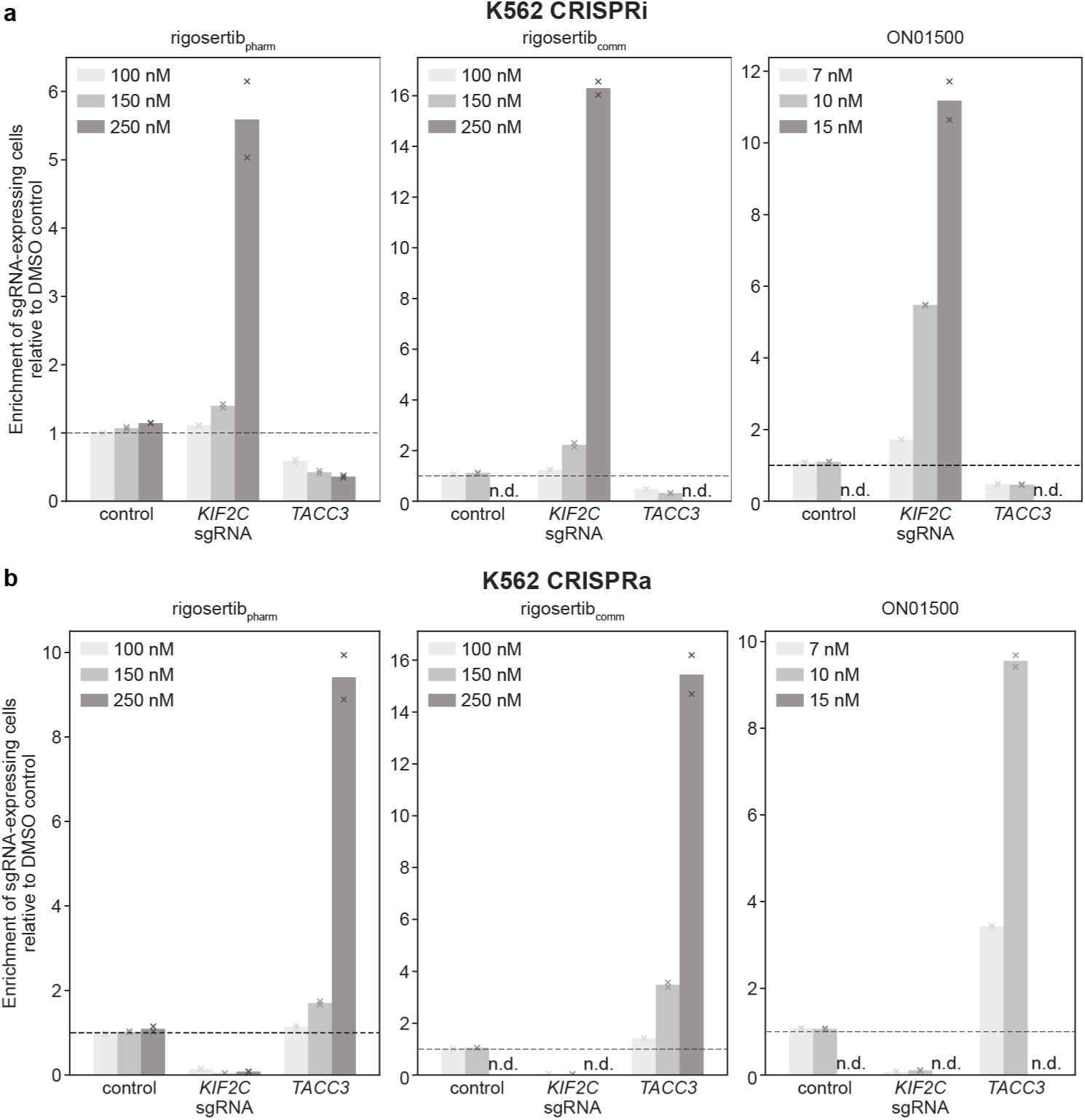
Internally controlled sensitivity assays to determine effects of *KIF2C* or *TACC3* knockdown or overexpression on sensitivity to rigosertib_pharm_, rigosertib_comm_, or ON01500. (a) CRISPRi drug sensitivity phenotypes for indicated sgRNAs. (b) CRISPRa drug sensitivity phenotypes for indicated sgRNAs. Enrichment is defined as ratio of sgRNA-positive cells to sgRNA-negative cells, normalized to the corresponding ratio after treatment with DMSO. n.d.: phenotype not determined because total counted cell numbers were <2,500. Data represent mean and individual measurements of replicate treatments (n=2).

### Pharmaceutical-grade rigosertib destabilizes microtubules in cells and in vitro

We next examined whether rigosertib_pharm_ affects microtubule dynamics in cells. Briefly, we performed time-lapse fluorescence microscopy on cells expressing the microtubule plus-end tracking protein EB3 fused to GFP to measure the dynamics of astral microtubules. In our original manuscript, we had used low doses of rigosertib and examined microtubule growth persistence in mitosis (as we found that rigosertib affected mitotic spindle assembly and resulted in a mitotic arrest). However, for ease of analysis, here we used a higher dose of rigosertib and examined microtubule growth rates in interphase, as drugs that bind to the colchicine site also affect microtubule growth rates (Jordan, 2002; Jordan and Wilson, 2004; Mohan et al., 2013; Stanton et al., 2011). We observed that microtubule growth speeds in cells were strongly affected by rigosertib_pharm_: treatment of cells with 2 µM rigosertib_pharm_ for 1 h reduced the growth speed of microtubules in cells 2.5-fold (Figure 2), demonstrating that rigosertib_pharm_ inhibits microtubule growth in cells.

**Figure 2.**
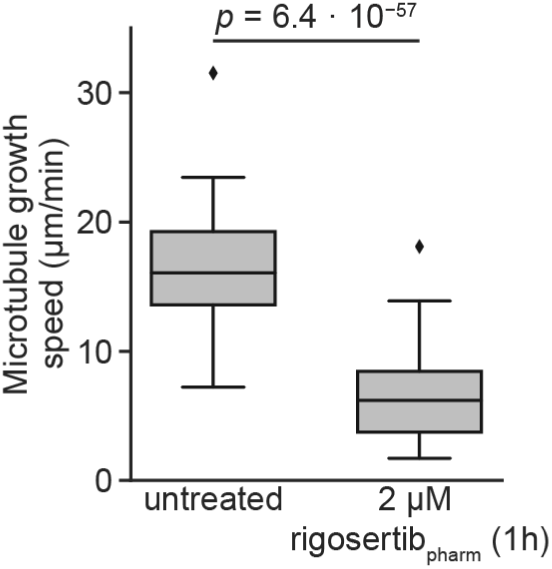
Rigosertib inhibits microtubule growth in cells. Microtubule growth speeds measured in untreated cells or cells treated with 2 µM rigosertib_pharm_ for 1 h. Untreated: n = 21; rigosertib: n = 29. Boxes denote IQR, central lines denote median values, whiskers denote lowest/highest datum within lower/higher quartile ± 1.5 IQR. Indicated *p*-value derived from a one-sided Mann-Whitney U-test.

We also assayed the effects of rigosertib_pharm_ on microtubule dynamics in vitro. To maximize sensitivity, we tracked the growth of individual microtubules reconstituted in vitro, in the presence of EB3, by time-lapse fluorescence microscopy. As we had observed in our original study, 10 µM rigosertib_pharm_ both reduced the growth speed of microtubules (Figure 3a) and increased the catastrophe frequency (Figure 3b). These data indicate that rigosertib_pharm_ can directly bind to tubulin and inhibit its polymerization. Our results disagree with the results from the bulk tubulin-polymerization assay shown in the manuscript by Reddy and co-workers. However, the assay used in our experiments is substantially more sensitive in detecting effects on microtubule growth (see Discussion). We also note that we observe these effects at 10 µM rigosertib, the lowest concentration tested in our experiments, in contrast to the inaccurate claims by Reddy and co-workers that we only observe in vitro microtubule destabilization at concentrations of 20 µM or higher. It is also important to note that many microtubule-destabilizing agents require substantially higher concentrations in vitro for robust microtubule-destabilizing activity as compared to cell culture (Panda et al., 1996; Jordan and Wilson, 2004; Mohan et al., 2013), possibly because these drugs accumulate over time in cells or because cellular factors modulate the effectiveness of microtubule-destabilizing agents. Therefore, the observed microtubule-destabilizing activity of rigosertib at < 10 µM in vitro is not unexpected, considering the high nanomolar concentrations required for cell killing. These results confirm that rigosertib_pharm_ directly destabilizes microtubules in vitro.

**Figure 3.**
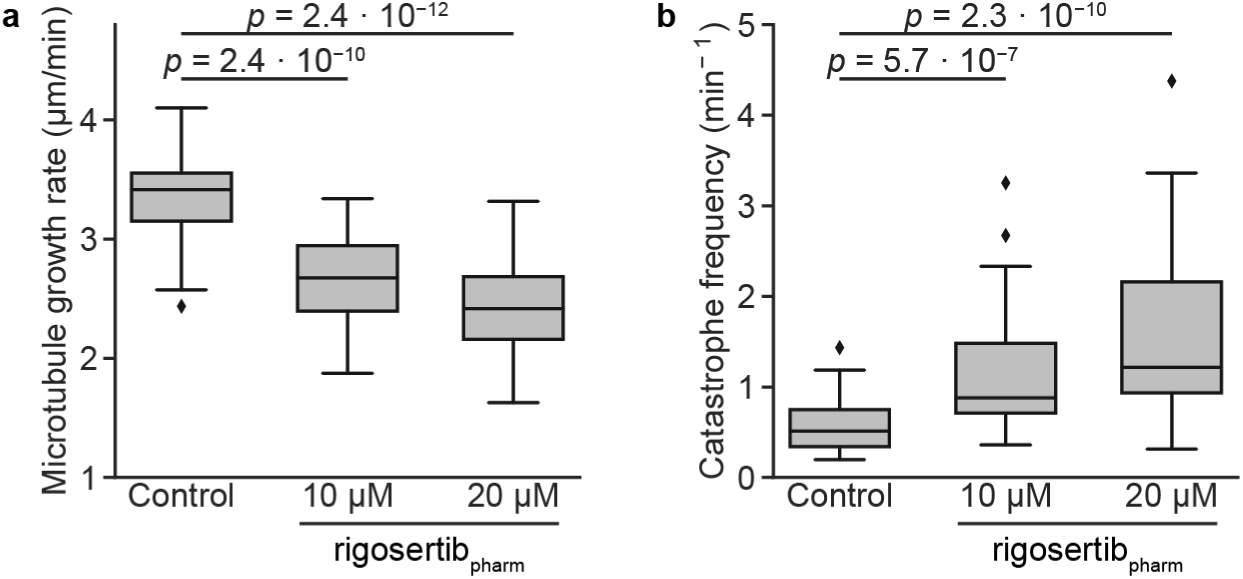
Rigosertib_pharm_ destabilizes microtubules in vitro. Quantification of **(a)** microtubule growth rate and **(b)** catastrophe frequency with 15 µM tubulin along with EB3 (20 nM) without or with 10 or 20 µM rigosertib_pharm_. n = 40 for each condition. Boxes denote IQR, central lines denote median values, whiskers denote lowest/highest datum within lower/higher quartile ± 1.5 IQR. Indicated *p*-values derived from one-sided Mann-Whitney U-tests.

### Expression of a mutant β-tubulin (L240F) protects against toxicity induced by pharmaceutical-grade rigosertib

We next evaluated if binding to tubulin is required for the cytotoxic activity of rigosertib_pharm_. In our original study, we had found that expression of a β-tubulin mutant with a mutation in the rigosertib binding pocket (L240F *TUBB*) conferred resistance to rigosertib. To determine if this mutant also conferred resistance to rigosertib_pharm_, we transduced K562 cells with a construct for expression of L240F *TUBB* from a constitutive SFFV promoter linked to mCherry via an internal ribosome entry site (IRES). We mixed transduced and wild-type K562 cells, exposed the mixture to rigosertib_pharm_ or ON01500, and measured the fraction of L240F *TUBB*-expressing cells over time as mCherry-positive cells by flow cytometry. Indeed, L240F *TUBB*-expressing cells enriched over wild-type cells upon treatment with both rigosertib_pharm_ and ON01500 (Fig. 4a), indicating that expression of L240F tubulin protects cells from toxicity induced by both compounds. Contrary to the claims of Reddy and co-workers that L240F *TUBB*-expressing cells undergo senescence and do not proliferate in the presence of rigosertib_pharm_, we found that L240F *TUBB*-expressing cells were actively proliferating in the presence of rigosertib_pharm_ at the same rate as DMSO-treated cells (Fig. 4b). These results demonstrate that expression of the mutant tubulin provides resistance to rigosertib_pharm_ and strongly suggest that tubulin binding by rigosertib is critical for its cytotoxic activity.

**Figure 4.**
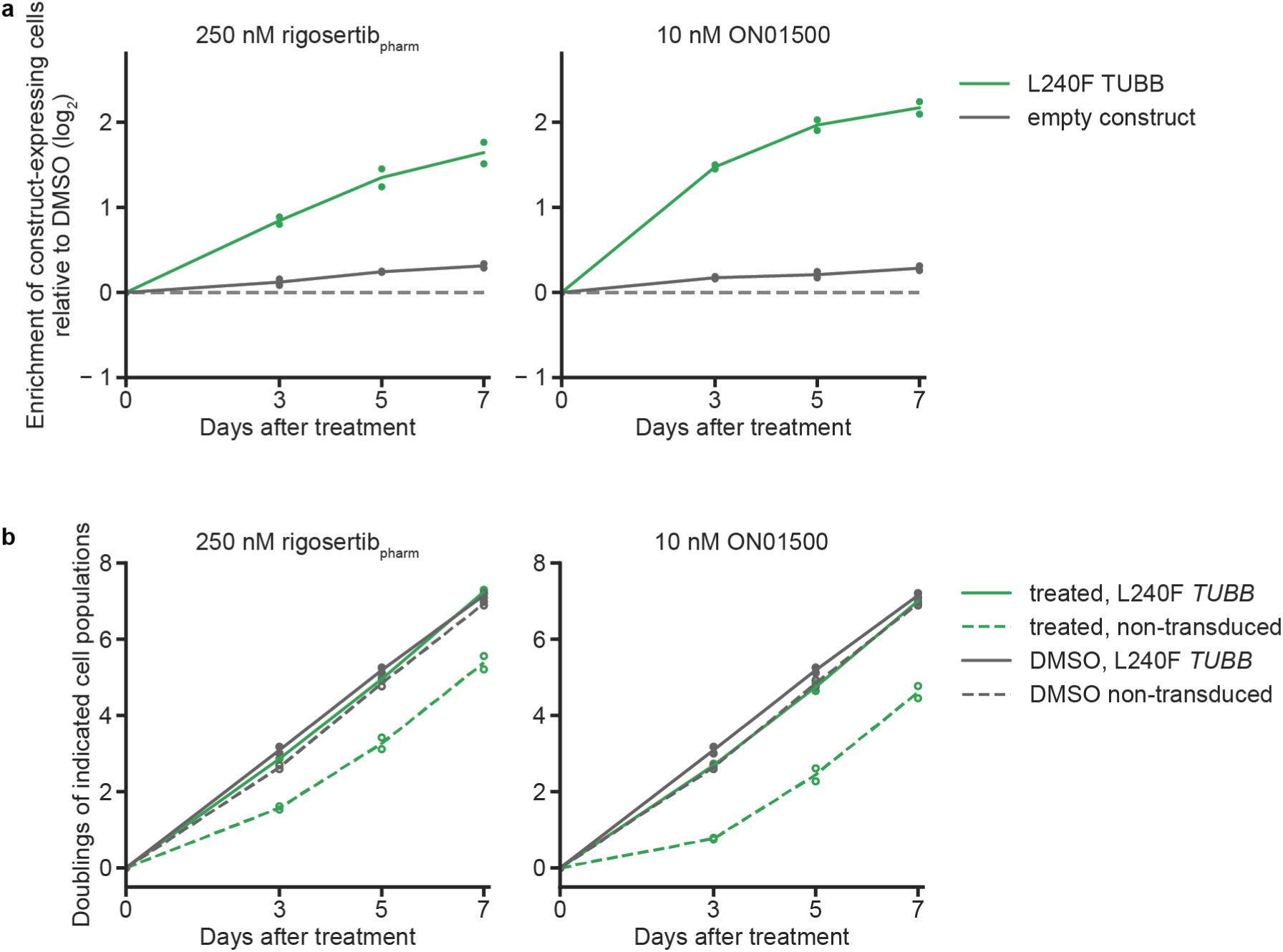
Expression of L240F *TUBB* confers resistance to rigosertib_pharm_. (**a**) log_2_ enrichment of K562 cells expressing L240F *TUBB* or an empty construct after treatment with rigosertib_pharm_ or ON01500 in internally controlled growth assays. Enrichment was measured as the ratio of mCherry-positive to mCherry-negative cells [e = fraction(mCh^+^) / fraction(mCh^−^)] by flow cytometry, calculated relative to the first time point. Relative enrichment for each time point was normalized to that of DMSO-treated control cells. Data represent mean and individual measurements of replicate treatments (n=2). (**b**) Cumulative cell doublings of L240F *TUBB*-transduced or non-transduced K562 subpopulations treated with rigosertib_pharm_ or ON01500. Cumulative doublings were calculated from measurements of cell numbers and the fractions of mCherry-positive (L240F *TUBB*-transduced) and mCherry-negative cells (non-transduced) in the population. Data represent mean and individual measurements of replicate treatments (n=2). Traces for DMSO-treated cells are identical in both panels in (**b**).

### Re-analysis of the crystal structure of tubulin complexed with rigosertib

In their manuscript, Reddy and co-workers suggested that our crystal structure of rigosertib-bound tubulin was more accurately represented by modeling ON01500 and a water molecule rather than rigosertib. To re-evaluate the structure, we refined models of either rigosertib or ON01500 against the deposited data and compared the resulting electron density maps at different contour levels (Figure 5a-c). We also calculated polder maps by omitting the ligands (a polder map is an omit map in which the bulk solvent around the omitted region is excluded; in this fashion, weak electron densities, which can be obscured by bulk solvent, may become visible). Modeling ON01500 and a water molecule indeed results in a good fit to the electron density (Figure 5b), but the polder maps show clear density that overlaps completely with rigosertib including the portion that distinguishes it from ON01500 (Figure 5c). Thus, based on the X-ray data it is not possible to unambiguously distinguish if our structures contain rigosertib, ON01500, or a mixture of the two compounds. Indeed, given the chemical similarity between ON01500 and rigosertib, the fact that the chemical differences are external to the main tubulin contacts, and the observation that both compounds alter microtubule stability in vitro and in cells, it seems likely that both compounds bind to this site. Regardless, we show that expression of the L240F *TUBB* mutant, which we had selected due to the proximity of the L240 residue to rigosertib in our original crystal structure, conferred resistance to rigosertib_pharm_, strongly supporting the conclusion that rigosertib binds to tubulin in the mode we described in our original manuscript.

**Figure 5.**
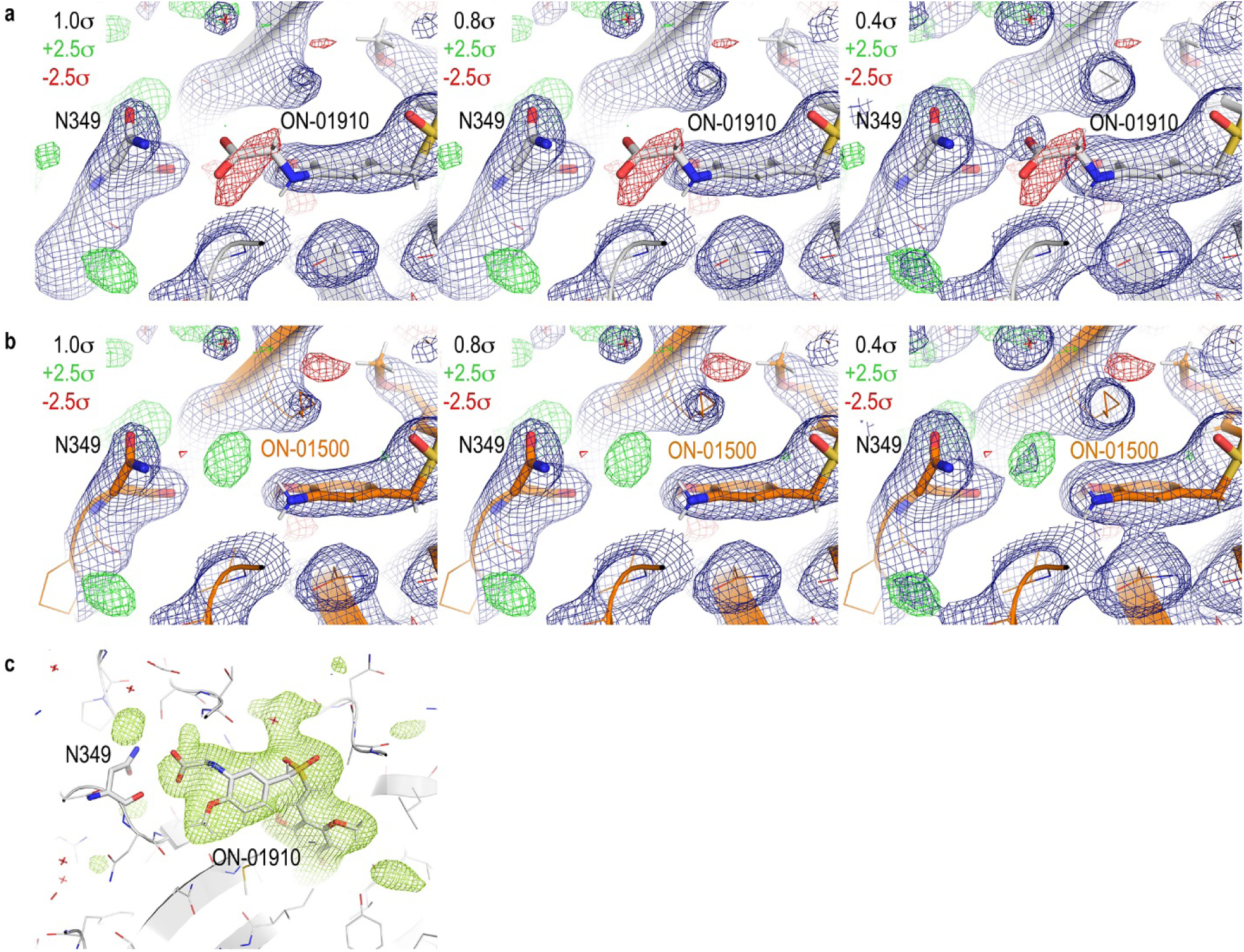
Reanalysis of the crystal structure of the tubulin-rigosertib complex. **(a)** Electron density of region in question after refinement of rigosertib against deposited data. 2*F*_o_–*F*_c_ (blue) and *F*_o_–*F*_c_ (green/red) is contoured at the indicated levels. **(b)** Electron density of region in question after refinement of ON01500 against deposited data. 2*F*_o_–*F*_c_ (blue) and *F*_o_–*F*_c_ (green/red) is contoured at the indicated levels. **(c)** Polder map around rigosertib contoured at 3.0 σ.

## Discussion

The broad goal of our original manuscript was to highlight the power of unbiased chemical-genetic screens to identify the mechanism of action of small molecules. We used rigosertib as a test case, and our screens directly pointed to microtubule destabilization as rigosertib’s mechanism of action, a hypothesis we confirmed using multiple orthogonal targeted assays. Although Reddy and co-workers agree that commercially obtained rigosertib kills cells by destabilizing microtubules, they raise the possibility that this microtubule-destabilizing activity was mediated by a contaminating impurity, ON01500 (Baker et al., 2019). They argue that pharmaceutical-grade rigosertib that lacks this impurity kills cells by inhibiting RAS signaling, PLK1 inhibition, and/or PI3K inhibition. Here, we re-evaluated the activity of pharmaceutical-grade rigosertib (>99.9% pure) as well as that of the potential impurity, ON01500. Similar to the work of Reddy and co-workers, we find that ON01500 is a potent microtubule-destabilizing agent. Importantly, we demonstrate that pharmaceutical-grade rigosertib also exhibits potent microtubule-destabilizing activity, with qualitatively indistinguishable behavior to the commercial rigosertib used for our original study in a series of different assays. Our data provide compelling evidence that rigosertib is a microtubule-destabilizing agent and that this activity is responsible for its cytotoxic activity.

On the surface, some of our results contradict those presented by Reddy and co-workers. Specifically, 1) we find that rigosertib inhibits microtubule growth in vitro, whereas Dr. Reddy’s team finds no effect of rigosertib on microtubule stability in vitro; 2) we find that expression of the resistant *TUBB* mutant specifically confers resistance to rigosertib, whereas Reddy and co-workers argue that this resistance is non-specific; and 3) we find that expression of the resistant *TUBB* mutant allows cells to proliferate in the presence of rigosertib, whereas Reddy and co-workers suggest that the cells do not proliferate and undergo senescence. Although these contradictions are difficult to reconcile without detailed knowledge of the protocols used to conduct their experiments, as discussed below our assays were designed to consistently provide higher specificity to detect the phenotypes in question. In addition, a key claim by Reddy and co-workers regarding the specificity of the rationally designed rigosertib-resistant *TUBB* mutant is contradicted by results published by independent investigators (Patterson et al., 2019).

### Effects of rigosertib on microtubules in vitro

We find using single-molecule fluorescence assays that pharmaceutical-grade rigosertib directly destabilizes microtubules in vitro at concentrations of 10 µM, the lowest concentration tested. By contrast, Reddy and co-workers find no effect of rigosertib on tubulin polymerization in bulk polymerization assays. We note that single-molecule fluorescence assays are far more sensitive than bulk tubulin polymerization assays, for which negative results are generally not interpretable. Indeed, several well-established microtubule-destabilizing agents including noscapine (Zhou et al., 2003) and griseofulvin (Panda et al., 2005) do not show an effect in bulk tubulin polymerization assays, but effects can be detected with more sensitive assays. These previous studies potentially explain why Reddy and co-workers did not detect rigosertib-induced microtubule destabilization in their bulk tubulin polymerization assays.

### Specificity of resistance conferred by the L240F *TUBB* mutant

We disagree on multiple grounds with the contention by Reddy and co-workers that the resistance conferred by the L240F *TUBB* mutant is non-specific. First, in our original study we demonstrated that expression of L240F *TUBB* provides resistance to rigosertib as well as ABT-751, a microtubule-destabilizing agent that binds in the same site on tubulin as rigosertib, but does not provide resistance to vinblastine, another microtubule-destabilizing agent that binds at a remote site on tubulin.

Second, Reddy and co-workers claim that the L240F *TUBB* mutant confers non-specific resistance because in their hands expression appeared to confer mild resistance to the PLK1 inhibitor BI2536. Patterson et al., however, recently published data from an essentially identical experiment and found that expression of the L240F *TUBB* mutant did not confer resistance to BI2536 (see Figure 7 in (Patterson et al., 2019)). Patterson et al. arrived at this experiment in a similar fashion as we did in our original study, by systematically characterizing the mechanism of action of a compound of interest, in this case the MTH1 inhibitor TH588. They found that TH588 synergizes with PLK1 inhibition in a manner that is independent of inhibition of MTH1, and through in-depth analysis found that TH588 destabilizes microtubules by binding in the same site as rigosertib. The L240F *TUBB* mutant indeed conferred resistance to TH588, but not to BI2536, which they used as a control (Patterson et al., 2019). Although this observation is published, Reddy and co-workers do not address the conflicting results, which are in support of our conclusions. We find it impossible to reconstruct what may have led to these conflicting results and thus choose not to speculate about the origins. Regardless, the combination of our data and those presented independently by Patterson et al. indicate that the L240F *TUBB* mutant confers resistance specifically to 3 inhibitors that all bind in the same site – rigosertib, ABT-751, and TH588 – but not to other cytotoxic agents including BI2536 as well as vinblastine, which also destabilizes microtubules but binds at a distal site. These results firmly establish the specificity of the L240F *TUBB* mutant.

### Extent of rigosertib resistance conferred by the L240F *TUBB* mutant

We had originally demonstrated in our manuscript that rigosertib-treated cells expressing the L240F *TUBB* mutant proliferated at the rate of DMSO-treated cells (Fig. S6F of (Jost et al., 2017)). In particular, in that figure we plotted the cumulative doubling <u>differences</u> compared to DMSO-treated cells, which were close to 0 for cells expressing the *TUBB* mutant. We repeated the same analyses with pharmaceutical-grade rigosertib and again found that rigosertib-treated cells expressing the *TUBB* mutant proliferated at the same rate as DMSO-treated control cells over the course of multiple days, whereas the addition of pharmaceutical-grade rigosertib induced a growth defect in cells that did not express the *TUBB* mutant. Thus, rigosertib-treated cells expressing L240F *TUBB* are clearly not senescent, as suggested by Reddy and co-workers, but are actively proliferating.

It is also important to clarify an inaccurate assumption made by Reddy and co-workers: the assumption that the L240F *TUBB* mutant should confer complete resistance to rigosertib at concentrations that are above the lethal level for wild-type cells. Indeed such complete resistance to rigosertib would be <u>unexpected</u> under these experimental conditions. In particular, rigosertib binding to tubulin subunits in microtubules inhibits growth and stimulates microtubule catastrophes; thus, any residual binding to microtubules even in the presence of the L240F *TUBB* variant would cause toxicity. There are at least three sources of such binding: the L240F *TUBB* variant likely retains some ability to bind rigosertib, rigosertib can still bind to alternative tubulin isoforms (cells express multiple tubulin genes), and finally, wild-type *TUBB* is still present in the cells as the L240F mutant is expressed in trans. Indeed, in our original work we had demonstrated that resistance to rigosertib is enhanced when endogenous *TUBB* is depleted by CRISPRi, but this important point is not acknowledged by Reddy and co-workers and in their experiments, wild-type *TUBB* is not depleted.

In retrospect there has been evidence for rigosertib’s microtubule-destabilizing activity in the literature since its first description, such as in the observation of multipolar spindles, a phenotype of low dose microtubule-destabilizing agents, in rigosertib-treated cells published by Reddy and co-workers in 2005 (Gumireddy et al., 2005). The multipolar spindle phenotype was attributed by Reddy and co-workers to PLK1 inhibition, but a substantial body of literature has now shown that PLK1 inhibition does not result in multipolar spindles (Lénárt et al., 2007; Steegmaier et al., 2007). Microtubule destabilization could certainly explain the anti-cancer activity of rigosertib, as other microtubule-destabilizing agents have long been mainstays of multiple chemotherapy regimens. As with any compound, a formal possibility is that the ultimate mechanism is mediated by a breakdown product, in which case the compound should perhaps more accurately be classified as a pro-drug. If that were the case for rigosertib, it would appear that such degradation would be inevitable under even rigorous experimental conditions and thus would be an essential aspect of rigosertib’s mechanism of action, as our results clearly suggest that pharmaceutical-grade rigosertib kills cancer cells by destabilizing microtubules.

More broadly, our re-evaluation further highlights the power of unbiased chemical-genetics to establish the mechanisms of action of small molecules even in the face of pleiotropy and chemical complexity. In our view, such approaches should ideally be employed for therapeutic candidates before the initiation of human trials to ensure that these candidates are deployed at maximum efficacy. Indeed, off-target activity of anti-cancer drugs appears to be more common than previously anticipated (Lin et al., 2019) and limits efficacy in targeted clinical trials, providing further motivation for the use of unbiased approaches to establish the in vivo targets for drugs that were developed through targeted assays.

## AUTHOR CONTRIBUTIONS

MJ, AR, AP, AA, MOS, MET, and JSW designed experiments. MJ, AR, and MET conducted experiments and analyzed data. AP and MOS analyzed the crystal structure. MJ, MET, and JSW wrote the manuscript. All authors edited the manuscript.

## ACKNOWLEDGEMENTS

We thank Onconova Therapeutics, specifically Dan Fox and Dr. V.J. Rajadhyaksha, for providing pharmaceutical-grade rigosertib and pure ON01500 for this study. This work was funded by the National Institutes of Health (NIH, grants P50 GM102706, U01 CA168370, R01 DA036858 to JSW; post-doctoral fellowship F32 GM116331 to MJ), the Swiss National Science Foundation (31003A_166608 to MOS), the European Research Council (Starting grant ERC-STG 677936-RNAREG to MET and Synergy grant 609822 MODELCELL to AA), a fellowship from the Dutch Cancer Society (KWF) to MET, NWO CW ECHO grant 711.015.005 to AA, NIH/NCI Pathway to Independence Award K99 CA204602 to LAG, and NIH/NCI Pathway to Independence Award K99 CA181494 and a Stand Up to Cancer Innovative Research Grant to MK. JSW is a Howard Hughes Medical Institute Investigator. MAH, LAG, MK, and JSW have filed a patent application related to CRISPRi and CRISPRa screening (PCT/US15/40449). MET, LAG, and JSW have filed a patent application for the SunTag technology (PCT/US2015/040439). JSW consults for and holds equity in KSQ Therapeutics, Maze Therapeutics, and Tenaya Therapeutics and is a venture partner at 5AM. MJ and MAH consult for Maze Therapeutics.

## STAR Methods

### CONTACT FOR REAGENT AND RESOURCE SHARING

Further information and requests for resources and reagents should be directed to and will be fulfilled by the Lead Contact, Jonathan S. Weissman.

### EXPERIMENTAL MODEL AND SUBJECT DETAILS

K562 cells were grown in RPMI 1640 (Gibco) with 25 mM HEPES, 2 mM L-glutamine, 2 g/L NaHCO_3_ and supplemented with 10% (v/v) fetal bovine serum (FBS), 100 units/mL penicillin, 100 µg/mL streptomycin, 2 mM L-glutamine. HEK293T cells were grown in Dulbecco’s modified eagle medium (DMEM, Gibco) with 25 mM D-glucose, 3.7 g/L NaHCO_3_, 4 mM L-glutamine and supplemented with with 10% (v/v) FBS, 100 units/mL penicillin, 100 µg/mL streptomycin. RPE1 cells were grown in DMEM:F12 (1:1) medium (Gibco) supplemented with 10% (v/v) FBS, 100 units/mL penicillin, and 100 µg/mL streptomycin. K562, and RPE-1 cells are derived from female patients/donors. All cell lines were grown at 37 °C.

## METHOD DETAILS

### Reagents

Pharmaceutical-grade rigosertib and ON01500 were obtained from Onconova through a material transfer agreement. Commercial rigosertib was obtained from SelleckChem.

### DNA transfections and virus production

Lentivirus was generated by transfecting HEK39T cells with standard packaging vectors using TransIT®-LT1 Transfection Reagent (Mirus Bio). Viral supernatant was harvested 2-3 days after transfection and filtered through 0.44 µm PVDF filters and/or frozen prior to transduction.

### Individual evaluation of sgRNA phenotypes

For individual evaluation and re-testing of sgRNA phenotypes, individually cloned sgRNA protospacers targeting *KIF2C* or *TACC3* or a non-targeting control protospacer (neg_ctrl-1) were used from our original study (Jost et al., 2017). The resulting sgRNA expression vectors were individually packaged into lentivirus and internally controlled growth assays to evaluate drug sensitivity phenotypes for each sgRNA were performed as described in our original study (Jost et al., 2017). Cells were transduced with sgRNA expression constructs at MOI < 1 (15 – 40% infected cells), treated with the corresponding drugs at approximately LD_60_ or DMSO 5 days after infection, and the fraction of sgRNA-expressing cells was measured 3 days and 5 days after treatment as BFP-positive cells by flow cytometry on an LSR-II (BD Biosciences). Specifically, for each treatment, 250,000 cells were seeded in one well of a 24-well plate for each population in duplicate in 500 µL complete RPMI containing the final desired drug concentration (day 0). The next day (day 1), 500 µL of fresh complete RPMI were added to dilute the drugs, and the subsequent day (day 2), 500 µL of the cell suspension were transferred to a new 24-well plate and again diluted with 500 µL of fresh complete RPMI. On day 3, both the fraction of sgRNA-expressing cells and cell density were measured by flow cytometry, and cells were split back to 250,000 cells in 1 mL of complete RPMI, or supplemented back to 1 mL complete RPMI if the total cell count was lower than 250,000 cells. The measurement was repeated on day 5, at which point the experiment was terminated.

### EB3-GFP tracking to measure microtubule growth speeds

RPE1 stably expressing dCas9-BFP-KRAB and EB3-GFP were seeded in 96-wells glass bottom dishes (Matriplate, Brooks). Immediately prior to imaging the medium was replaced by Leibovitz’s L-15 (Gibco) CO_2_-independent medium supplemented with or without the indicated concentration of rigosertib. The cells were imaged using a Yokogawa CSU-X1 spinning disk confocal attached to an inverted Nikon TI microscope with Nikon Perfect Focus system, 100× NA 1.49 objective, an Andor iXon Ultra 897 EM-CCD camera, and Micro-Manager software (Edelstein et al., 2014). 50 images were acquired for each movie at 1 s time interval in a single z-section through the middle of the cell. To measure microtubule growth speeds, kymographs were created along growing microtubules. Microtubule growth speeds were calculated based on the slope of lines in the kymographs.

### In vitro microtubule polymerization assays

To monitor the direct effects of rigosertib on microtubule dynamics, in vitro assays (as described previously (Doodhi et al., 2016) and in our original manuscript (Jost et al., 2017)) were performed with reaction mixtures in MRB80 buffer containing tubulin (15 µM), Rhodamine-tubulin (0.5 µM) when indicated, methyl cellulose (0.1%), KCl (50 mM), k-casein (0.5 mg/ml), GTP (1 mM), oxygen scavenging system (20 mM glucose, 200 μg/ml catalase, 400 μg/ml glucose-oxidase, 4 mM DTT), mCherry-EB3 (20 nM) and with different concentrations of rigosertib. Movies were acquired in total internal reflection fluorescence (TIRF) microscopy mode using a Nikon Eclipse Ti-E (Nikon) microscope supplemented with the perfect focus system (PFS) (Nikon), equipped with a Nikon CFI Apo TIRF 100x 1.49 N.A. oil objective (Nikon) and a photometrics CoolSNAP HQ2 CCD (Roper Scientific) camera with triple-band TIRF polychroic ZT405/488/561rpc (Chroma) and triple-band laser emission filter ZET405/488/561m (Chroma), mounted in the metal cube (Chroma, 91032) together with emission filter wheel Lambda 10-3 (Sutter instruments) with ET460/50m, ET525/50m and ET630/75m emission filters (Chroma). Vortran Stradus 488 nm (150 mW) and Cobolt Jive 561 nm (100 mW) lasers were used for excitation (the laser launch was part of ILas system (Roper Scientific France/ PICT-IBiSA, Institut Curie)) at a laser power of 6 with an exposure time of 100 ms. Images were acquired with MetaMorph 7.7 software (Molecular Devices) at 63 nm per 1 pixel. Kymographs were generated by ImageJ using the KymoResliceWide plugin. Two independent assays were performed for each condition to collect the reported data.

### L240F TUBB rescue assay

The rescue assay used constructs for stable expression of L240F *TUBB* or an HA tag (empty vector control) from a constitutive SFFV promoter, linked to mCherry via an IRES. Note that in our original manuscript, the constructs were expressed from an inducible TRE3G promoter, but similar results were obtained here in a simpler fashion with the constitutive SFFV promoter. The constructs were individually packaged into lentivirus and transduced into K562 CRISPRi cells at a multiplicity of infection ≤1 (30-60% infected cells). To measure effects on drug sensitivity, cells were treated with drugs or DMSO 5 days after infection and the fraction of *TUBB*-expressing cells was measured 3 days after treatment and then every 2 days as the fraction of mCherry-positive cells by flow cytometry on an LSR-II flow cytometer (BD Biosciences). Specifically, for each treatment, 250,000 cells were seeded in one well of a 24-well plate for each cell population in duplicate in 500 µL complete RPMI containing the final desired drug concentration (day 0). The next day (day 1), 500 µL of fresh complete RPMI were added to dilute the drugs, and the subsequent day (day 2), 500 µL of the cell suspension were transferred to a new 24-well plate and again diluted with 500 µL of fresh complete RPMI. On day 3, both the fraction of *TUBB*-expressing cells and cell density were measured by flow cytometry, and cells were split back to 250,000 cells in 1 mL of complete RPMI, or supplemented back to 1 mL complete RPMI if the total cell count was lower than 250,000 cells. This procedure was repeated on days 5 and 7, at which point the experiment was terminated.

## QUANTIFICATION AND STATISTICAL ANALYSIS

For all experiments, details of quantification and statistical methods used are described in the corresponding figure legends or results sections. The methods used to quantify microtubule growth properties in cells and *in vitro* are described above.

